# Community rescue in experimental phytoplankton communities facing severe herbicide pollution

**DOI:** 10.1101/467324

**Authors:** Fugère V., Hébert M.-P, Costa N.B., Xu C.C.Y., Barrett R.D.H., Beisner B.E., Bell G., Fussmann G.F., Shapiro B.J., Yargeau V., Gonzalez A.

## Abstract

Evolutionary rescue occurs when adaptation prevents local extinction in deteriorating environments. Laboratory experiments with microorganisms have shown that the likelihood of evolutionary rescue is greatest in large populations that have previously experienced sublethal doses of stress. To assess this result in natural communities, we conducted a mesocosm experiment with semi-natural phytoplankton communities exposed to glyphosate, a widely used herbicide. We tested whether community biomass and pre-exposure to sublethal stress would facilitate community rescue after severe contamination. Exposure to sublethal stress, but not community biomass, facilitated rescue significantly–even though it led to biodiversity loss. Furthermore, glyphosate had modest effects on community composition, suggesting that community resistance to glyphosate was primarily driven by changes in resistance within taxa, not by community turnover. Our results expand the scope of community evolutionary rescue theory to complex ecosystems and confirm that prior stress exposure is a key predictor of rescue.

---

Human-induced global change has led to unprecedented rates of population extirpation and species extinction^1–3^, a ‘biodiversity crisis’ that can have profound impacts on ecosystem functions and services^4,5^. However, rapid evolution could potentially mitigate biodiversity loss in degraded environments via the process of ‘evolutionary rescue’^6,7^. Evolutionary rescue (ER) occurs when stress-resistant genotypes spread to high frequency in a population facing severe environmental deterioration, thus allowing a demographic recovery of the population while changing its genetic composition^8^. Assuming sufficient adaptive variation for stress resistance (supplied by pre-existing variation or new mutations), two key factors that influence the incidence of ER in degraded environments are population size prior to environmental degradation and pre-exposure to sublethal doses of stress^9^. The former influences the risk of stochastic extinction while the population experiences a decline in abundance at the onset of stress^10–13^. The latter creates selection that increases the frequency of stress-resistant genotypes in the population, thus allowing it to withstand more severe doses of stress thereafter^11,14,15^.

Most empirical studies of ER have used microorganisms in laboratory environments, such that the incidence of ER in nature remains controversial^9,16,17^. Moreover, ER experiments have traditionally focused on single species because early theory involved single-species models^8^. Recent theory also predicts ER in communities exposed to stress^18,19^. In line with this theory, one laboratory experiment exposed multiple co-occurring species of soil microbes to a lethal dose of a novel stressor (the herbicide, Dalapon) and observed the simultaneous ER of multiple taxa, which allowed overall community abundance to recover under severely-degraded conditions^20^. This experiment suggested the possibility of ‘community rescue’, defined as the recovery or maintenance of an aggregate community property such as biomass under conditions that, without adaptation, are normally lethal to all constituent populations of the community. The likelihood of community rescue appears to depend on some of the same factors that predict ER in single-species experiments, e.g. community abundance (summed across populations/species) and the history of stress (prior exposure) of the community^20^.

We extended this research and assessed, for the first time, community rescue in complex communities under semi-natural conditions, using plankton in pond mesocosms as a model system (Fig. 1a). We used the pesticide glyphosate to induce severe herbicide pollution, which is known to have toxic effects on several species of phytoplankton^21–24^. Glyphosate is the most widely-used pesticide worldwide, with an applied tonnage rising sharply and continuously since the development of glyphosate-resistant crops in the early 1990s^25–27^. Traces of glyphosate in the environment have led to concerns over potential health and ecotoxicological impacts^28–32^. Moreover, many plant species have evolved glyphosate-resistance in recent years^33,34^, creating weed management problems^35^, but also suggesting that communities could potentially adapt rapidly to this contaminant and undergo ER when exposed to high doses^36^.

**Figure 1.**
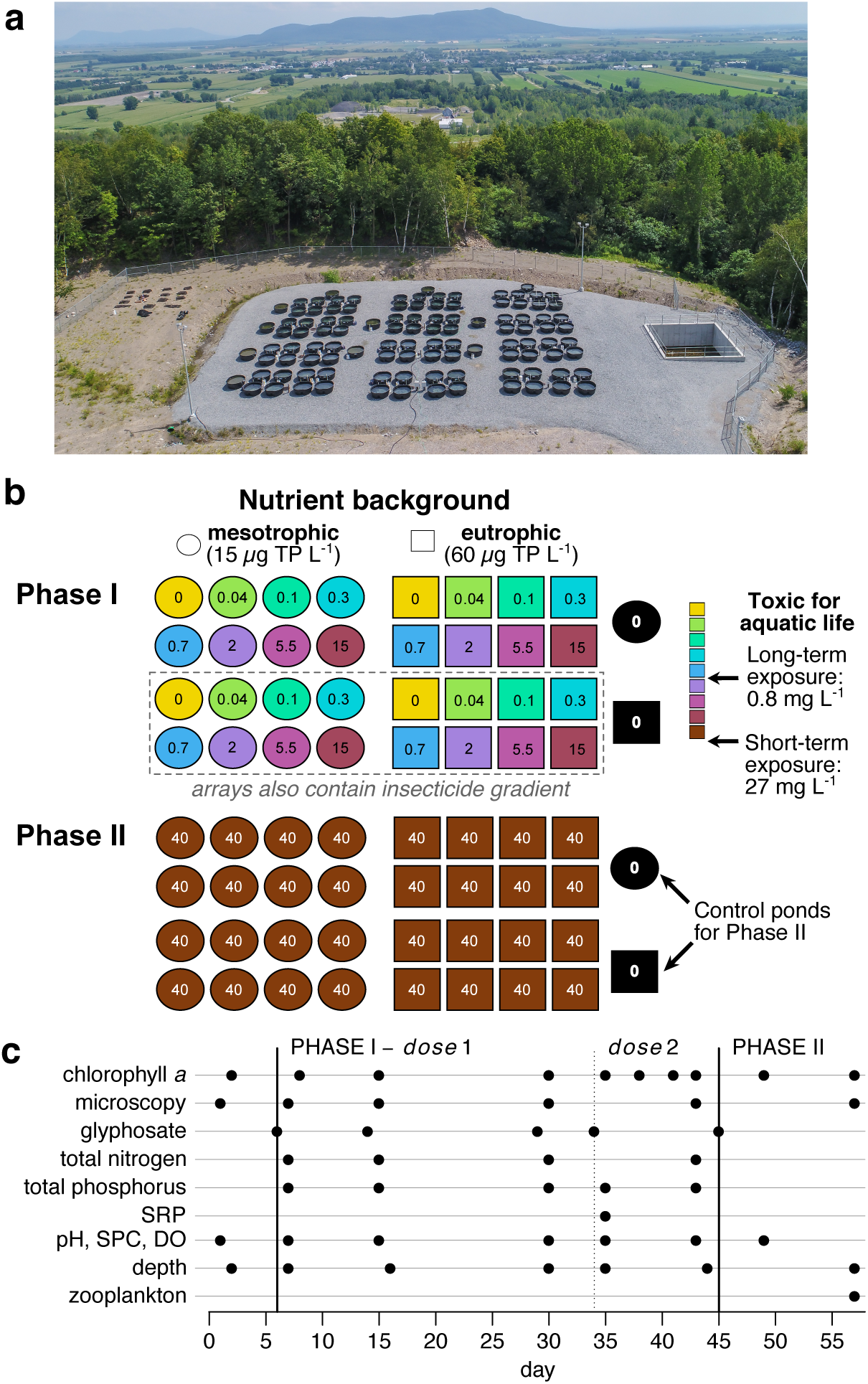
Experimental site, design, and timeline. **(a)** Aerial photograph of the Large Experimental Array of Ponds facility at Gault Nature Reserve, located near an area of intensive agriculture. **(b)** Schematic representation of experimental treatments. Colours and numbers within symbols indicate target glyphosate concentrations after application of one dose. The nutrient treatment was a press treatment maintained with biweekly nutrient addition. The glyphosate treatment involved, in Phase I, two pulse applications (doses) of Roundup ranging in concentration from 0-15 mg/L of glyphosate acid, and in Phase II, one dose of 40 mg/L in all experimental ponds. Yellow and black ponds are pesticide-free in Phase I, while yellow ponds (but not black ponds) receive the lethal dose in Phase II. **(c)** Timeline of the experiment. Symbols indicate measurement dates for variables listed on the left. Temperature was also recorded in all ponds with automated sensors. Thick vertical lines indicate the beginning of Phase I and II, while the dotted line indicate the second dose of Phase I. TP = total phosphorus; SRP = soluble reactive phosphorus; SPC = specific conductance; DO = dissolved oxygen.

We conducted a community rescue experiment with 34 pond mesocosms inoculated with a diverse phytoplankton community originating from a pristine lake in Southern Québec. The lake is located on a mountain within a forested protected area, itself surrounded by a region of intensive agriculture of glyphosate-resistant corn and soy where traces of glyphosate have been detected in nearly all lower-lying water bodies monitored by local authorities^37^. We tested whether this naïve phytoplankton community could be rescued from severe glyphosate pollution, and if so, whether rescue would be facilitated by higher community biomass and pre-exposure to sublethal stress, as in the laboratory community rescue experiment described above^20^. The experiment had two phases (Fig. 1b). In Phase I, we imposed divergent selection for 40 days, manipulating community biomass (with a press nutrient treatment) and pre-exposure to sublethal stress (with two pulse applications of Roundup–a commercial glyphosate formulation–varying in concentration). Then, in Phase II, all ponds (excepting two controls) were exposed to a dose of Roundup expected to be lethal after short-term exposure. Throughout the experiment, we tracked phytoplankton biomass (chlorophyll *a* concentration), community composition (genus-level biovolume), and water chemistry, including glyphosate and nutrient concentrations (Fig. 1c). We also measured zooplankton density at the end of the experiment. Community biomass at the end of Phase II indicates the potential of a community to maintain its productivity in a severely-degraded (normally lethal) environment and is our measure of community rescue, which we relate to the two factors manipulated in Phase I (community biomass and prior stress exposure).

## Results

At the start of the experiment (day 2), one week after the first nutrient application, high-nutrient ponds had a greater phytoplankton biomass than low-nutrient ponds (GAM, nutrient effect: *p* = 0.003; Fig. 2a,b). This positive effect of nutrient enrichment on phytoplankton biomass remained significant throughout Phase I of the experiment (GAM, nutrient effect: *p* = 0.007; Fig. 2a,c-e). In contrast, and as expected, ponds assigned to different glyphosate treatments did not differ in phytoplankton biomass prior to the first pesticide pulse (GAM, effect of ‘future glyphosate dose’: *p* = 0.393; Fig. 2a,b). The two pulse applications of glyphosate during Phase I of the experiment then had a strong, time-dependent effect on biomass (GAM, interaction effect of time and glyphosate concentration: *p* < 0.0001; Fig. 2a,c-e). When we applied the first glyphosate pulse (day 6), the pesticide had a negative, dose-dependent impact on phytoplankton biomass, reducing chlorophyll *a* concentration to < 1 μg/L in ponds receiving the highest dose (Fig. 2a,c). However, even the most impacted communities recovered quickly, and effects of glyphosate on phytoplankton biomass were no longer evident by day 15–even if glyphosate concentration remained constant during this period (Fig. 2a; Fig. S1a,b).

**Figure 2.**
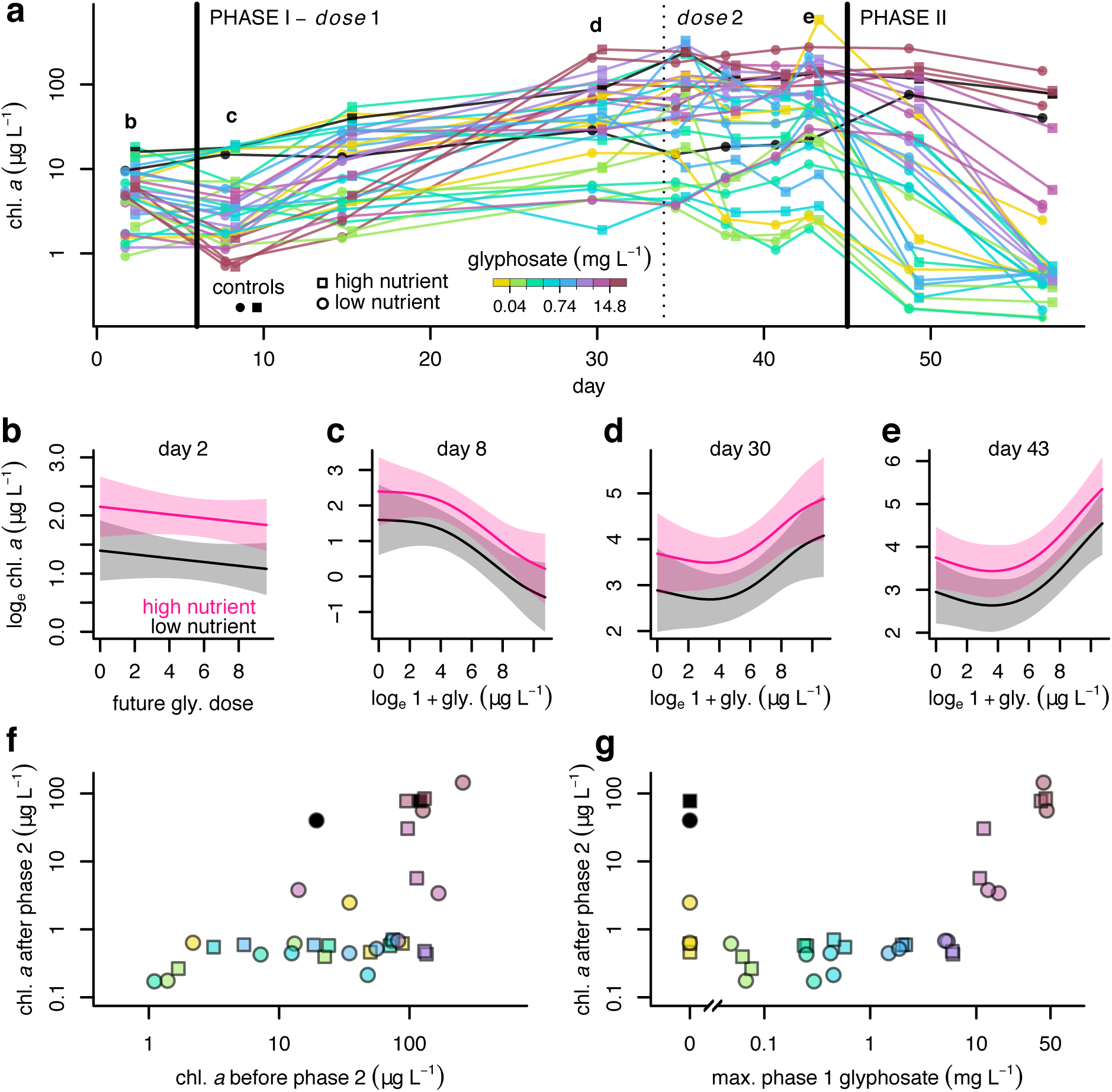
Phytoplankton biomass dynamics during the experiment. (a) Time series of chlorophyll a concentration (a proxy for phytoplankton biomass) in all ponds over the course of the experiment. Symbols and colours indicate nutrient and glyphosate treatments, respectively. Black lines/symbols are control ponds for Phase II. **(b-e)** Results of additive mixed models predicting chlorophyll a concentration from measured glyphosate concentration and nutrient treatment. Model results are shown for various key time points of Phase I. Shaded polygons illustrate 95 % confidence intervals. **(f-g)** Chlorophyll a concentration at the end of Phase II as a function of chlorophyll a at the end of Phase I (g) or maximum recorded glyphosate concentration during Phase I (g). chl. = chlorophyll; gly. = glyphosate.

Then, from day 15 to 30, before a second dose was applied, phytoplankton biomass increased steeply in the high-glyphosate ponds, and the effect of glyphosate had reversed to a positive, dose-dependent impact on phytoplankton biomass (Fig. 2a,d). We then applied a second dose of glyphosate on day 34, which led to significantly higher in-pond glyphosate concentrations than what we had targeted (Fig. S1a,b). This was due to the lack of degradation of the first pulse as well as evaporation and a gradual decline in water level during Phase I (Fig. S2a). Despite glyphosate concentration exceeding 30 mg/L in some ponds, this second, unintentionally more severe dose did not have a negative effect on biomass–rather, the glyphosate-biomass relationship remained positive after the second dose (Fig. 2e), and chlorophyll *a* concentration reached values > 100 μg/L in all high-glyphosate ponds by the end of Phase I (Fig. 2a).

We attribute the longer-term, fertilizing effect of Roundup during Phase I to the nutrient content of the glyphosate molecule (8.3 % nitrogen and 18.3 % phosphorus; other compounds in Roundup such as the surfactant polyethoxylated tallow amine also contain nutrients). Bioavailable nutrients could be released and potentially assimilated by organisms upon degradation of the pesticide; for example, inorganic phosphorus-containing compounds are among the main degradation products of glyphosate^38,39^. Although we did not note obvious degradation of glyphosate when measuring in-pond concentration over multiple days after the first pulse application (Fig. S1a-b), concentration of soluble reactive phosphorus (SRP; mostly orthophosphate) was significantly higher in ponds receiving the highest glyphosate doses (Fig. S3), indicating that at least partial glyphosate degradation and bioavailable P release had occurred. The nutrient content of Roundup also led to a strong, dose-dependent increase in total nitrogen (TN) and total phosphorus (TP) concentrations during Phase I (Fig. S1c-d). This effect was markedly stronger than our nutrient treatment, which reached the target concentrations of 15 and 60 μg/L TP in control ponds only (Fig. S1d). In high-glyphosate ponds, TP concentrations exceeded 1 mg/L, although most of this phosphorus could remain biologically unavailable. In contrast, the glyphosate and nutrient treatments had little influence on other physicochemical parameters. Depth and temperature varied over time but not across mesocosms (Fig. S2a,b). Mean specific conductance increased slightly over Phase I (from 91 to 116 μS/cm), indicative of solute accumulation in the mesocosms due to evaporation (Fig. S2c). Dissolved oxygen concentration tracked changes in phytoplankton biomass and was negatively affected by the first glyphosate pulse in the ponds exposed to the highest dose (Fig. S2d). pH was mostly stable over time, although the highest glyphosate doses temporarily lowered pH by < 1 unit (Fig. S2e).

The lack of biomass decline following the second glyphosate dose of Phase I suggests that community resistance was increased by the first dose. In Phase II of the experiment, when all experimental communities were contaminated with a severe dose of glyphosate expected to be lethal (target in-pond concentration = 40 mg/L), biomass indeed collapsed in most communities (Fig. 2a). However, some communities remained as productive as the control communities, indicating community rescue. Community rescue (biomass at the end of Phase II) was unrelated to both community biomass before degradation (GAM, effect of Phase I chlorophyll *a*: *p* = 0.377; Fig. 2f) and to nutrient treatment (GAM, nutrient effect: *p* = 0.355; squares vs. circles in Fig. 2f,g). In contrast, the extent of glyphosate exposure during Phase I was a very strong predictor of rescue (GAM, effect of Phase I glyphosate: *p* < 0.0001; Fig. 2g), confirming that glyphosate-exposed communities acquired greater glyphosate resistance during Phase I. Biomass collapse in communities that did not rescue also decreased dissolved oxygen concentration (Fig S2d), while specific conductance and pH respectively increased and decreased in all ponds that received the lethal dose irrespective of the response of their phytoplankton community (Fig. S2c,e). No obvious change in phytoplankton biomass or water chemistry was noted for the two control ponds during Phase II (Fig. 2a,f-g; Fig. S2), confirming that seasonal changes in temperature or irradiance cannot explain biomass collapse in glyphosate-treated ponds which did not rescue.

Interestingly, because glyphosate added during Phase I did not degrade significantly, some high-glyphosate communities that retained functionality (high biomass) in Phase II were also those that were exposed to the most extreme concentrations. For example, in two high-glyphosate ponds, Phase II glyphosate concentration exceeded 80 mg/L (Fig. S1a). However, we also noted significant variability in Phase II glyphosate concentration that could not be accounted for by residual glyphosate from previous applications (Fig. S1a,b). For example, a few high-nutrient ponds had much lower concentrations than expected (Fig. S1a). This variability in Phase II glyphosate concentration is likely due to measurement error as opposed to a failure to apply the same amount of Roundup in all ponds. For example, it seems very unlikely that we would have consistently applied less Roundup to high than low-nutrient ponds (and indeed, nutrient treatment had no effect on Phase II phytoplankton biomass). Moreover, the biomass response of all ponds within a given glyphosate treatment was very consistent (Fig. 2g). We nonetheless tested for an effect of measured Phase II glyphosate concentration on Phase II phytoplankton biomass and found a positive relationship (the opposite of one might expect) driven entirely by rescue in high-glyphosate ponds (Fig. S4; see also the last paragraph of this section).

Although biomass recovered in ponds receiving a high dose of glyphosate in Phase I, phytoplankton diversity did not. Indeed, in the subset of ponds for which we collected composition data, we observed a gradual loss of diversity in high-glyphosate ponds over the course of Phase I (Fig. 3a,d). At the end of Phase I, glyphosate concentration had a weak but significant negative effect on both genus number (GAM, effect of glyphosate: *p* = 0.0447; Fig. 3b) and alpha diversity measured as the effective number of genera (GAM, effect of glyphosate: *p* = 0.0143; Fig. 3e). The nutrient treatment had a significant negative impact on the effective number of genera (GAM nutrient effect: *p* = 0.0162; Fig. 3e) but not genus number (GAM nutrient effect: *p* = 0.505; Fig. 3b). At the end of Phase II, both rescued and collapsed communities had generally lower diversity than control communities (Fig. 3c,f).

**Figure 3.**
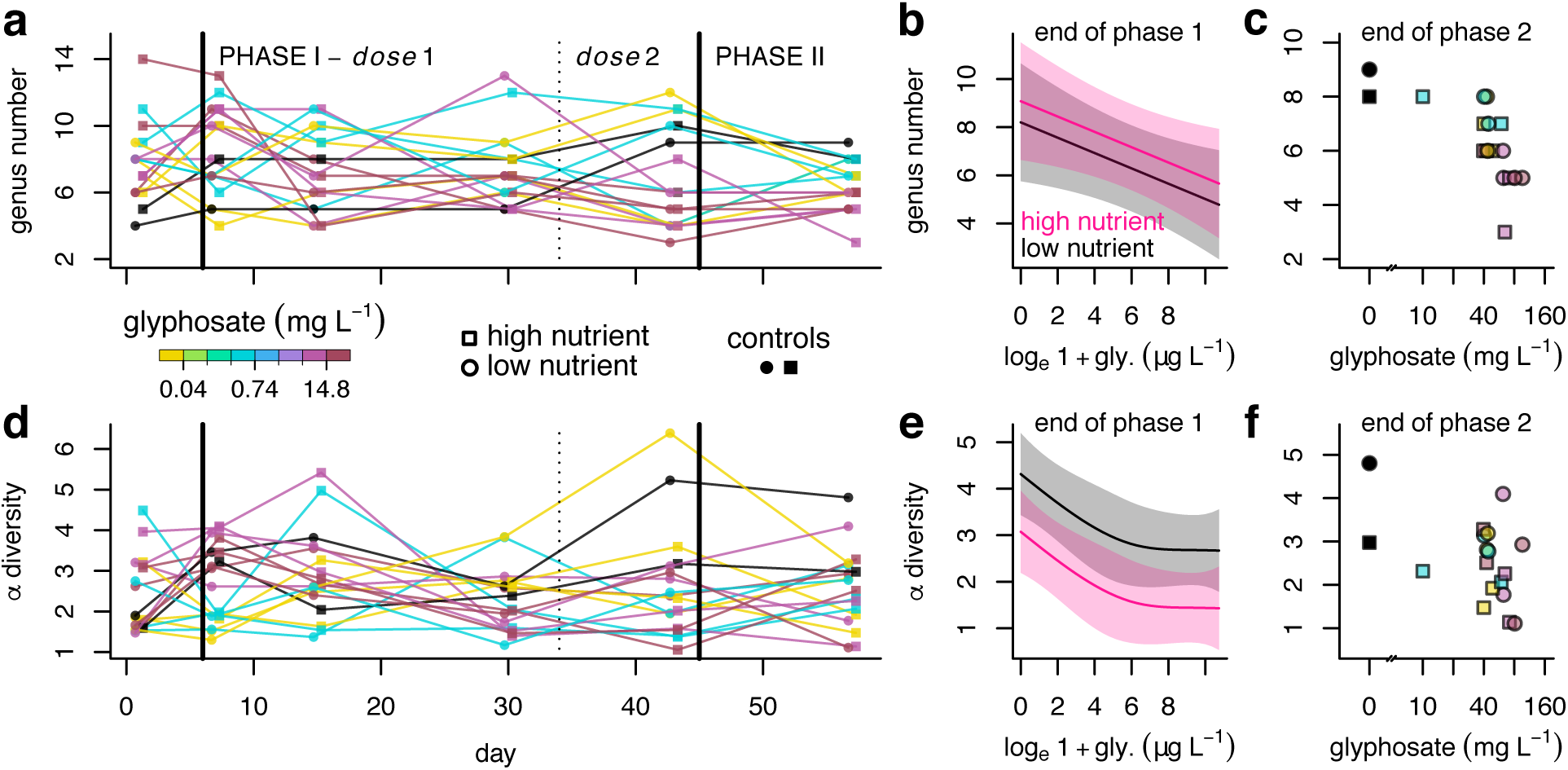
Effect of glyphosate on phytoplankton biodiversity. **(a, d)** Time series of rarefied richness (a) and *α* diversity (effective number of genera; d) in the subset of ponds for which we collected composition data. Symbols and colours are as in Fig. 2. **(b, e)** Results of additive mixed model predicting richness (b) or diversity (e) at the end of Phase I as a function of glyphosate concentration and nutrient treatment. **(c, f)** Richness (c) and diversity (f) of communities at the end of Phase II in relation to measured glyphosate concentration at the onset of Phase II. Colours and symbols indicate glyphosate and nutrient treatments as in (a). gly. = glyphosate.

In spite of this overall negative effect on diversity, glyphosate exposure had a modest influence on community composition because a few taxa (*Selenastrum*, *Ankistrodesmus*, *Desmodesmus*, and *Chlorella)* were highly-dominant in all ponds. When comparing community composition at the beginning vs. end of Phase I using the Bray-Curtis dissimilarity index, we noted that all ponds diverged from their starting composition regardless of their nutrient or glyphosate treatment (Fig. 4a). Dissimilarity at the end of Phase I, i.e. the extent of community divergence over the first 44 days of the experiment, was not significantly related to glyphosate exposure (GAM glyphosate effect: *p* = 0.731; Fig. 4b) nor nutrient treatment (GAM nutrient effect: *p* = 0.193; Fig. 4b). Community synchrony (*η*), expected to be more negative (asynchronous) in high-glyphosate ponds if the herbicide induced significant genus sorting^40^, was indeed slightly more negative in high-glyphosate ponds, but only for the high-nutrient treatment (GAM, effect of glyphosate on *η* in high-nutrient ponds: *p* = 0.0102; effect of glyphosate in low-nutrient ponds: *p* = 0.8832; Fig. 4c). Moreover, synchrony values were all close to zero, indicating that dynamics of different genera were mostly uncorrelated, even in high-glyphosate, high-nutrient ponds. Community composition was also weakly related to glyphosate exposure during Phase I (Fig. 4d). Indeed, although composition was initially similar across ponds (Fig. 4d, open symbols), communities diverged in directions not predicted by their experimental treatments (Fig. 4d, full symbols). At the end of Phase I, high-glyphosate communities showed marked differences in composition, while one unexposed community had a composition similar to 3 high-glyphosate ponds. This suggests that various ‘routes to resistance’ were possible in high-glyphosate ponds during Phase I, and/or that stochasticity and ecological drift had a stronger influence on community reassembly than environmental forcing by the glyphosate gradient. Furthermore, not only was glyphosate treatment a poor predictor of community composition (Fig. S5a,b), but community composition at the end of Phase I was itself a poor predictor of rescue during Phase II (Fig. S5c,d).

**Figure 4.**
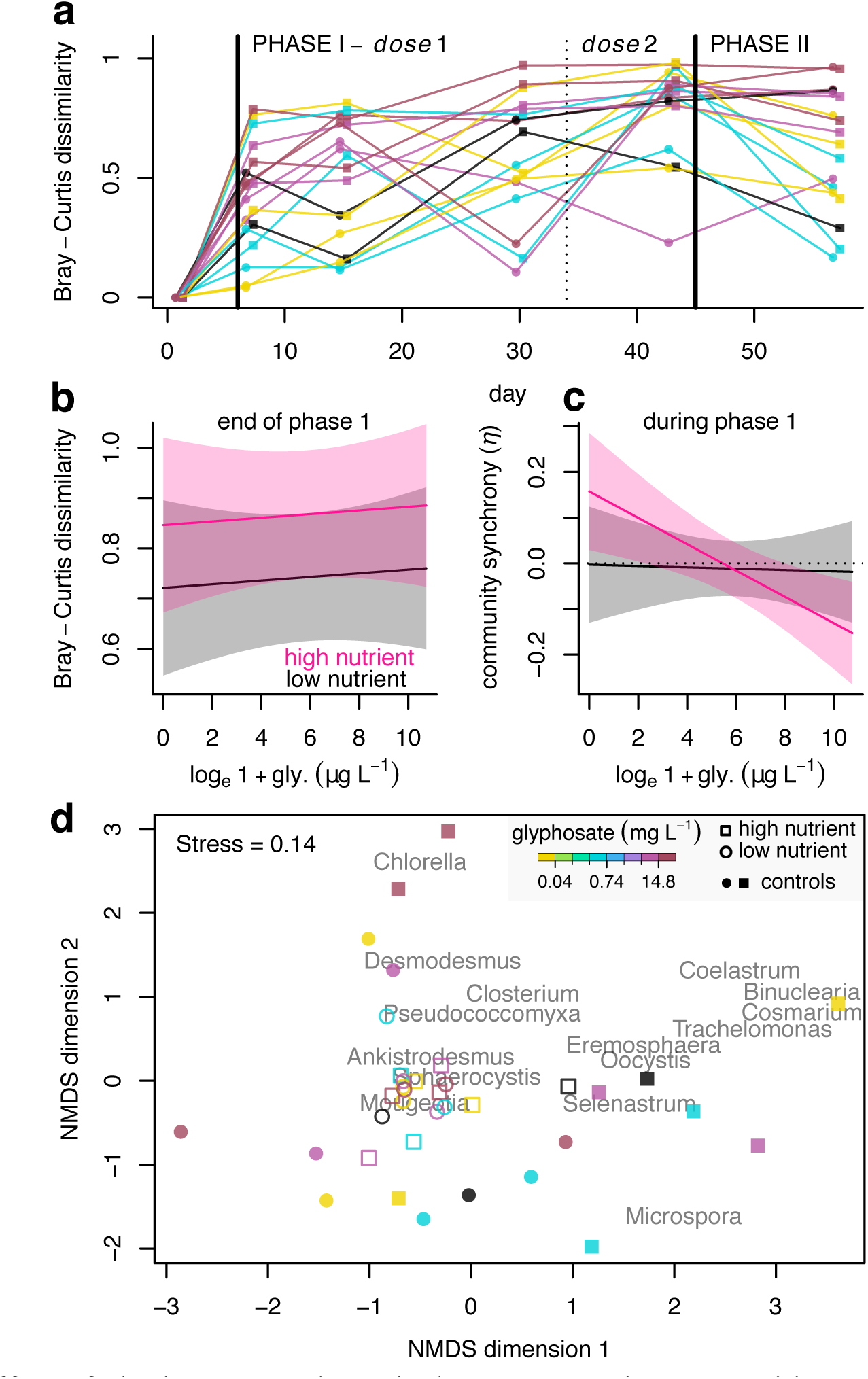
Effect of glyphosate on phytoplankton community composition. **(a)** Time series of Bray-Curtis dissimilarity of each pond relative to its starting composition. Higher values indicate greater community divergence over the course of the experiment. Symbols and colours are as in Fig. 2. **(b, c)** Results of additive mixed models predicting Bray-Curtis dissimilarity at the end of Phase I (b) or community synchrony (*η*) during Phase I (c) as a function of glyphosate concentration and nutrient treatment. For the synchrony index, more negative values indicate more asynchronous dynamics, while a value of zero indicates independent taxon fluctuations. **(d)** Non-metric multidimensional scaling (NMDS) representation of community composition at the beginning (open symbols) and end (full symbols) of Phase I. The position in two-dimensional space of the fifteen most abundant taxa is also shown. gly. = glyphosate.

To determine which properties of communities best predicted their likelihood of rescue in Phase II, we conducted two analyses in which stress exposure, biomass, diversity, and composition variables were all included as predictors of final phytoplankton biomass at the end of Phase II, in the 16 ponds for which data were available for all variables. We also included final crustacean zooplankton density as a predictor, as zooplankton grazing could have aggravated the collapse of phytoplankton biomass in naïve ponds. In a regression tree analysis, we found that glyphosate exposure in Phase I was the only variable necessary to distinguish rescued from collapsed communities; a threshold exposure concentration of 0.578 mg/L during Phase I determined final biomass at the end of Phase II (Fig. 5a). Then, when fitting and comparing independent GAMs with one of thirteen community properties as the predictor variable and biomass at the end of the experiment as the response, we found that glyphosate concentration at the end of Phase I was by far the best predictor of rescue (Fig. 5b). Zooplankton density was not a good predictor of rescue (Fig. 5b). Furthermore, the relationship between phytoplankton biomass and zooplankton density was positive, indicating weak top-down control of phytoplankton by zooplankton (Fig. S6). This (weak) positive relationship suggests that phytoplankton rescue influenced zooplankton density in Phase II rather than the opposite pathway of zooplankton grazing influencing phytoplankton rescue.

**Figure 5.**
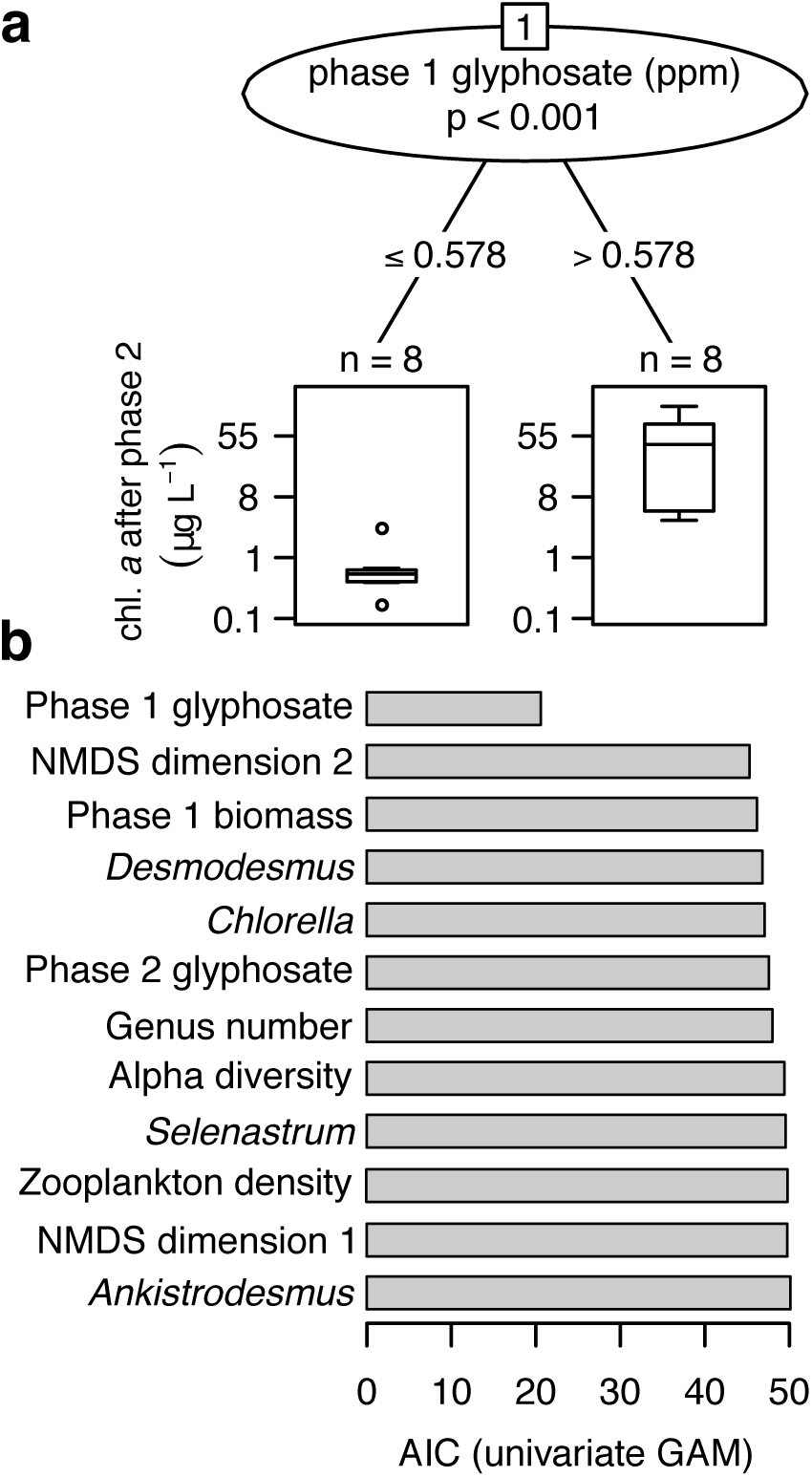
Predictors of community rescue. (a) Regression tree predicting phytoplankton biomass at the end of Phase II as a function of various community properties. Results (*p* value) of a permutation test of a correlation between the response and the one significant predictor (glyphosate exposure during Phase I) is indicated. (b) Model fit (AIC) of univariate generalized additive models (GAMs) with phytoplankton biomass at the end of Phase II as the response variable and one of the community properties used in (a) as the predictor variable. A lower AIC indicates better fit. Genus names represent relative biovolumes of a given taxon. chl. = chlorophyll; ppm = parts per million (mg/L).

## Discussion

Our results indicate that exposure to high doses of Roundup increases phytoplankton community resistance and prevents biomass collapse when the same communities are subsequently contaminated by a much higher concentration of glyphosate. This result is consistent with laboratory microcosm studies finding an influence of prior exposure on the likelihood of rescue^14,20^. Various processes could contribute to increased glyphosate resistance in the communities that remained productive in Phase II. In controlled experiments with single species^10,41^, adaptation can be inferred from a U-shaped demographic trajectory at the onset of stress. Indeed, a switch from negative to positive population growth in a constant (highly-stressful) environment is indicative of trait change, i.e. an increase in mean individual stress resistance within the population. Both phenotypic plasticity^42^ and genetic adaptation (from standing variation or from *de novo* mutations) can contribute to increased population-level stress resistance. However, in a multi-species experiment such as the one that we describe here, species sorting and compensatory dynamics could also increase stress resistance at the community level if taxa that are originally resistant to the stressor become relatively more abundant. That is, community rescue could involve both ecological and evolutionary processes, with selection and sorting of adaptive variation operating at both interspecific and intraspecific levels. These various mechanisms have also been discussed in the ecotoxicological literature on ‘stress-induced community tolerance’^43,44^, but in the context of community responses to multiple unrelated stressors.

We suggest that our results indicate a greater role for increased glyphosate resistance within taxa than for sorting, at least at the genus level (the taxonomic resolution of our biovolume data). Glyphosate treatment only induced weak sorting; the same genera could dominate control (glyphosate-susceptible) and exposed (glyphosate-resistant) ponds at the end of Phase I (see also^45^). Furthermore, the only common feature of glyphosate-resistant communities that remained productive in Phase II was their history of glyphosate exposure in Phase I. Neither community biomass nor composition predicted rescue; nor did the relative biovolume of taxa common in (some) resistant communities. Other forms of rescue such as demographic and genetic rescue^9^ can be ruled out as well, as we used closed communities of abundant microorganisms. Therefore, we hypothesize that community rescue in this experiment was principally driven by evolutionary and/or plastic rescue, which could be determined with follow-up genomic analyses. One key target of selection in the genome could be the 5-enolpyruvylshikimate-3-phosphate synthase (EPSPS) gene, the enzyme targeted by glyphosate and the locus of adaptation in most glyphosate-resistant weed species^36,46^. Molecular analyses will also help distinguish clonal selection within species (an evolutionary process) from species selection within genera (an ecological process), and thus overcome one important limitation of our community analyses based on genus-level microscopy data^47^.

Our results also highlight the dual effect of glyphosate on a naïve lake phytoplankton community: herbicidal, at first, but fertilizing over a longer period. Importantly, negative effects on biomass and diversity were only observed at the highest experimental doses (> 2 mg/L). Such concentrations exceed by orders of magnitude concentrations typically measured in water bodies in agricultural areas, which are generally in the ng to μg/L range^28,30^ (although these low concentrations could in part be due to the rapid degradation of glyphosate in water). Moreover, we used Roundup, reputed to be even more toxic than pure glyphosate due to its surfactant^21,48,49^, and still recorded modest toxicity for both phyto- and zooplankton. Thus, in lakes with a plankton composition similar to our source community, runoff of glyphosate from agricultural fields will unlikely cause a significant loss of plankton biodiversity and biomass. However, the longer-term, fertilizing effect of Roundup on phytoplankton biomass was stronger than its initial toxic effect, and even the lowest doses in the μg/L range caused an increase in water nutrient concentrations. Other experimental studies have observed this fertilizing effect and have attributed it to the nutrient content of the herbicide^22,45,50,51^. In some phytoplankton species, the glyphosate molecule itself can be used as a resource even in the absence of microbial breakdown of glyphosate into simpler compounds^52^. Furthermore, all nutrients contained in commercial formulations of glyphosate applied to fields constitute a nutrient input that persists in the environment even after the herbicide degrades (unlike ecotoxicological effects, which eventually vanish once degradation is complete). In some areas of intensive culture of glyphosate-resistant crops, glyphosate application now constitutes a substantial source of anthropogenic phosphorus comparable in magnitude to other inputs that have been previously regulated^27^. Thus, a key environmental impact of glyphosate pollution might be via its effect on nutrient loading^22,51,53,54^, an issue that warrants further investigation given the extensive usage of this pesticide.

Our results extend one key finding from laboratory microcosm studies of ER to larger, more complex ecosystems: pre-exposure to sublethal stress permits community persistence in a severely-degraded environment that is otherwise lethal to naïve communities. Remarkably, communities selected in a glyphosate-rich environment for a few weeks only could remain productive when later facing a very high concentration of glyphosate (96 mg/L in the most contaminated pond). Our zooplankton data also suggests that rescue in primary producers could then sustain a viable consumer community in some severely-contaminated ponds. Nonetheless, the loss and recovery of biomass in Phase I that increased community resistance came at the expense of diversity, as glyphosate-resistant communities at the end of the experiment had 30-60% fewer genera than uncontaminated ponds. This loss of diversity suggests a cost of community rescue analogous to the demographic costs of adaptation at the population level^16,55^, which can reduce genetic diversity. One key avenue for future research will be to determine whether the loss of intra- and interspecific variation induced by rescue from one stressor influences the likelihood of rescue from another stressor^56–58^, to better define the limits of community rescue in human-dominated landscapes where multiple stressors often co-occur. Finally, although the prediction that the history of stress exposure predicts ER held true, the lack of an influence of community biomass on rescue in this experiment contrasts with results from microcosm studies^20^. Our approach demonstrates the value of testing ER theory with complex communities under more natural conditions. Evidence of ER in nature is accumulating^59–62^–the next challenge will be to determine which constituents of impacted communities can undergo rescue and whether they can sustain the recovery of ecosystem functions and services in degraded environments.

## Methods

### Experimental design

The experiment was conducted at the ‘Large Experimental Array of Ponds’ facility at McGill University’s Gault Nature Reserve in Québec, Canada (45°32′N, 73°08′W). This facility comprises > 100 mesocosms (1136 L Rubbermaid plastic tanks) that can be filled with water and planktonic organisms piped down from a lake (Lac Hertel) located 1 km upstream of the facility (Fig. 1a). Lac Hertel has a fully forested (and protected) watershed with no history of agriculture, and thus its community should be naïve to glyphosate. All mesocosms were filled on May 11^th^, 2016 with unfiltered lake water. Biweekly water changes of 10 % total mesocosm volume (with lake water and organisms) were performed until the experiment commenced. Major terrestrial inputs (pollen, leaves) were removed periodically with a leaf skimmer. Our 34-pond experiment then ran from August 17^th^ (day 1) to October 12^th^ (day 57), after which all mesocosm water was pumped into a sewage system that outflows into a large retention basin. Two months later, after glyphosate had degraded to a low concentration considered safe for aquatic life^63^ and for human consumption^64^, the water was released in a field outside of the protected area.

Fig. 1b illustrates our experimental design. In Phase I of the experiment (day 1-44), we manipulated community biomass and pre-exposure to sublethal stress. Then, Phase II (day 45-57) of the experiment represented our rescue assay, when all ponds (excepting two controls) were exposed to a high dose of Roundup expected to be lethal (see below). We manipulated community biomass in Phase I via a nutrient treatment, attributing 17 ponds to a ‘mesotrophic’ (low nutrient) treatment with a target total phosphorus (TP) concentration of 15 μg/L (similar to Lac Hertel), and 17 ponds to a ‘eutrophic’ (high nutrient) treatment with a target TP concentration of 60 μg/L (Fig. 1b). We prepared a concentrated nutrient solution of KNO_3_ (107.66 g/L), KH_2_PO_4_ (2.17 g/L), and K2HPO4 (2.82 g/L) with the same N:P molar ratio (33:1) as Lac Hertel in August 2016. Every two weeks for eight weeks, 5 or 20 ml of that stock solution were applied to low and high-nutrient ponds, respectively. The first nutrient addition took place on August 10^th^, one week before sampling started, to ensure that phytoplankton communities would have passed their exponential growth phase when applying the first pesticide pulse.

The glyphosate treatment of Phase I involved two pulses of Roundup Super Concentrate (Monsanto, St-Louis, MO, USA), applied on days 6 and 34. We used Roundup rather than pure glyphosate salt because local agricultural fields are sprayed with commercial formulations of glyphosate, not with the pure compound. Importantly, we used this herbicide as a generic stressor to induce environmental degradation; the precise mechanism of toxicity was not the focus of our study. Between mesocosms, Roundup doses varied in their target concentration (0-15 mg/L of glyphosate acid, the active ingredient in Roundup); a total of eight concentrations were used, separated by equal intervals on a logarithmic scale to cover a broad gradient (Fig. 1b; Phase I). Some doses used were greater than the Canadian aquatic toxicity criterion (environmental concentrations considered safe for aquatic life) for long-term glyphosate exposure, but the range of concentrations used falls below the criterion for short-term exposure^63^. These toxicity criteria are based on ecotoxicological assays with phytoplankton, plants, invertebrates, fish, and amphibians. The glyphosate gradient was repeated four times; twice at each nutrient level (totaling 32 ponds; Fig. 1b). We also included one additional pond at each nutrient level without pesticide application (shown in black in Fig. 1b) to serve as controls for Phase II; thus, there were 6 control (glyphosate-free) ponds in Phase I (3 of each nutrient level), but two control ponds for Phase II. Roundup was added to the mesocosms to reach the target concentrations, assuming a mean pond volume of 1000 L. Based on existing literature^50,65,66^, we expected glyphosate to degrade quickly before the second application and thus, both doses were expected to result in the same in-pond concentration.

Phase II began on day 45, when all ponds excepting two controls were treated with Roundup to reach a target in-pond concentration of 40 mg/L. This concentration, which exceeds the Canadian aquatic toxicity criterion for short-term exposure by 13 mg/L^63^, reduced phytoplankton biomasses to a very low level (< 1 μg/L) in a laboratory pilot experiment with water samples from the mesocosms. Community biomass at the end of Phase II (day 57), namely the capacity of a community to remain productive under severely deteriorated conditions that are normally lethal, was our measure of community rescue. Because the 34 ponds used in this study were also part of a larger (ecotoxicological) experiment with multiple agricultural stressors, two of the glyphosate gradients of Phase 1 (one at each nutrient level) also received a gradient of imidacloprid, a neonicotinoid insecticide. This insecticide gradient had no detectable effect on any of the response variables that we measured (see supplementary results in SI Appendix). Thus, both glyphosate gradients for each nutrient treatment were grouped and considered replicates.

### Sampling

The sampling schedule for each response variable is shown in Fig. 1c. All sampling equipment were thoroughly washed and dried between sampling occasions. Mesocosm water was sampled with integrated samplers made from 2.5 cm diameter PVC tubing. Samples were collected at 5 random locations in the upper 35 cm of the water column and combined in a 1 L dark Nalgene bottle, previously triple-washed with pond water. Each pond had a dedicated sampler and bottle to minimize cross-pond contamination. While sampling, bottles were kept in coolers and then transferred to an on-site laboratory. The 1 L samples were used to measure nutrient concentrations and phytoplankton biomass and composition (glyphosate samples were collected separately; see below). To estimate phytoplankton biomass, 50 ml was poured into a dark microcentrifuge tube. Chlorophyll *a* concentration, a proxy for phytoplankton biomass, was then determined fluorometrically with a FluoroProbe (bbe Moldaenke, Schwentinental, Germany). The FluoroProbe determines both total phytoplankton biomass (pigment concentration) and the biomass of four major groups that differ in their pigment coloration and fluorescence: green algae (chlorophytes), golden/brown algae (diatoms, chrysophytes, and dinoflagellates), blue-green algae (cyanobacteria), and cryptophytes.

To measure phytoplankton community composition at a finer taxonomic resolution in a subset of ponds (all four ponds receiving glyphosate dose 1 (controls), 4, 7 or 8), we preserved 45 ml samples with Lugol’s iodine solution for later microscopic enumeration. Samples were identified to genus level using the Utermöhl method^67^. Subsamples were sedimented in a 10 ml settling chamber and then screened using an inverted phase contrast microscope (Zeiss, Germany). A minimum of 200 cells and 10 fields were counted at both 100x and 400x magnification, to include both large and small cells. Ten fields at 40x magnification were also counted to identify large colonies. Colony number was multiplied by a genus-specific average number of cells per colony and then added to the cell count at higher magnification. Counts were converted to biovolume using a genus-specific mean cell volume obtained from a trait database for phytoplankton genera of Southern Québec (B.E. Beisner, unpublished data). Missing values for some taxa were obtained from a larger, published database^68^ accessed through the R package ‘phytotraitr’ (available from: https://github.com/andrewdolman/phytotraitr), using the median value reported for a given genus. For three (rare) taxa missing from this database, we used the value of a morphologically similar, closely related genus.

For nutrient concentrations, we retained 40 ml whole-water samples in acid-washed glass tubes, in duplicate each for total nitrogen (TN) and total phosphorus (TP). Samples were refrigerated until processed in the GRIL analytical laboratory at the Université du Québec à Montréal. Samples for TN were analyzed with a continuous flow analyzer (OI Analytical, College Station, TX, USA) using an alkaline persulfate digestion method, coupled with a cadmium reactor, following a standard protocol^69^. Phosphorus concentration was determined spectrophotometrically by the molybdenum blue method after persulfate digestion^70^. Pond TN and TP concentrations were estimated as the mean of the two duplicates. On day 36 of the experiment, one day after applying the second glyphosate dose, we measured TP and soluble reactive phosphorus (SRP) in 16 ponds (8 glyphosate doses × two nutrient treatments–in the two arrays without insecticide), to determine whether glyphosate applications increased SRP concentration. SRP was measured with the same protocol as TP but water samples were pre-filtered with 0.45 μm syringe filters to exclude particulate phosphorus.

To measure in-pond glyphosate concentration and validate that we established the target gradient, 1 L water samples were collected in clear plastic bottles immediately after applying Roundup. Samples were acidified to a pH < 3 with sulfuric acid and frozen until analysis. Samples were collected in all ponds after each application of Roundup, as well as in a subset of ponds (dose 1, 4, and 8; i.e. 0, 0.3, and 15 mg/L) 8 and 23 days after the first dose, to measure the rate of glyphosate degradation in our mesocosms. We also collected a sample of lake water to confirm that it had no glyphosate. Glyphosate concentration was later determined in the Department of Chemical Engineering at McGill University with liquid chromatography heated electrospray ionization tandem mass spectrometry using an Accela 600-Orbitrap LTQ XL (Thermo Scientific, Waltham, MA, USA). Acquisition was conducted in full scan mode (50-300m/z) at high resolution (FTMS=30 000m/Dz), with an ion trap used to perform targeted data acquisition for the product ion spectra (MS2) and generate identification fragments. The limits of detection and quantification of the method were 1.23 and 4.06 μg/L, respectively. Data were analyzed with Xcalibur 2.1.0 (Thermo Scientific).

Water pH, dissolved oxygen, and specific conductance were measured *in situ* in each mesocosm with a hand-held probe (YSI Inc., Yellow Springs, OH, USA) placed in the volumetric center of the pond. Measurements were taken at sunrise and sunset; the mean of both measurements was used to quantify the daily average. Depth in the center of the pond was recorded with a meter stick; we only measured depth in glyphosate-free ponds as little variation was observed across the array. Water temperature was recorded every 15 mins over the course of the experiment with HOBO pendant autonomous temperature data loggers (Onset, Bourne, MA, USA) deployed in all ponds. Finally, we also collected zooplankton samples at the end of the experiment. A total of 2 L of water collected with the integrated samplers at 10 random locations were combined and filtered with a 64 μm sieve. Zooplankton were anesthetized using carbonated water and then preserved in 95% ethanol to a final concentration of 75 % ethanol. Abundance and density of crustaceans (cladocerans and copepods) were determined microscopically.

### Statistical analyses

All analyses were conducted in R version 3.5.0^71^. Our analyses only included green algae because FluoroProbe data indicated that this group contributed 98.6 % of phytoplankton biomass when considering all ponds and sampling dates together. Rare golden/brown algae were detected at the onset of the experiment but went extinct quickly in all ponds irrespective of nutrient and glyphosate treatments. Other groups (e.g., cyanobacteria and cryptophytes) were exceedingly rare, with pigment concentrations comparable to the limit of detection of the FluoroProbe (< 0.1 μg/L; which is what we measured in distilled water).

Time series of chlorophyll *a* concentration (log-transformed) in Phase I were modelled using generalized additive mixed models (GAMs) fitted with the function ‘gam’ in the R package ‘mgcv’^72^. We used GAMs for most analyses to account for the non-linearity of many relationships, even when variables were log-transformed. To confirm that ponds from different glyphosate treatments did not initially differ in biomass, we first tested for an effect of nutrient treatment (a binary factor) and ‘future glyphosate dose’ (a smooth term corresponding to the log-transformed glyphosate treatment assigned to a given pond) on chlorophyll *a* on day 2, before the first glyphosate dose was applied. We then modelled chlorophyll *a* on all sampling occasions of Phase I as a function of nutrient treatment, time (a smooth term), glyphosate concentration measured in the pond (log-transformed; a smooth term), and ‘pond’ (a random effect). We fitted various models including only the nutrient effect, only the glyphosate effect, and/or both effects and all possible two-way interactions. The best model was selected using Akaike information criterion (AIC). This model had the following R syntax: chlorophyll ~ nutrient + s(date, glyphosate) + s(site, bs=‘re’). This model required a glyphosate concentration for all sampling occasions. Because we found no evidence of glyphosate degradation after the first pulse (see Results), glyphosate concentration in ponds that we did not sample on any given date was assumed to correspond to the concentration when the pond was sampled last (i.e. after a Roundup addition). To test the hypothesis that community biomass and pre-exposure to sublethal stress influence the likelihood of community rescue, we fitted a GAM with chlorophyll *a* at the end of Phase II as the response variable and nutrient treatment (a factor) and chlorophyll *a* and glyphosate concentration at the end of Phase I as predictors (two smooth terms). The three continuous variables were log-transformed. We only modelled Phase II chlorophyll *a* in ponds that received the lethal dose.

We then conducted a number of diversity and community composition analyses in the subset of ponds with genus-level biovolume data. Genus number and alpha diversity (effective number of genera^73^) were calculated for all ponds and time points. We used GAMs to test for an effect of glyphosate concentration and nutrient treatment on these two variables, on the last time point of Phase I. Diversity at the end of Phase II was also examined but no statistical test was performed since all ponds received the same glyphosate dose. Divergence in community composition (relative biovolume of each genus) over the course of the experiment was quantified with the Bray-Curtis dissimilarity index. For each pond, we calculated dissimilarity at each time point relative to initial composition on day 2. We also quantified community synchrony during Phase I (between day 2 and day 44), to determine whether glyphosate exposure led to asynchronous (compensatory) dynamics of individual genera. We estimated synchrony (*η*) with the R package ‘codyn’^74^, whereby *η* is the average correlation between the biovolume of each genus and the total biovolume of all other genera in the community^40^. An *η* value of 1 indicates perfect synchrony (all taxa fluctuate in sync), a value of -1 indicates perfect asynchrony among taxa (with biovolume remaining constant), and a value close to zero indicates independent fluctuations among genera. We then tested whether glyphosate exposure influenced community divergence and community synchrony by fitting GAMs with either dissimilarity at the beginning vs. end of Phase I (divergence) or *η* (synchrony) as the response, and with nutrient treatment, glyphosate concentration at the end of Phase I (log-transformed; a smooth term), and their interaction as predictors.

To visualize divergence in community composition during Phase I of the experiment, we constructed non-metric multidimensional scaling (NMDS) representations of community composition in two dimensions, including data from day 2 (before treatments) and day 44 (end of Phase I). NMDS analysis was performed with the ‘metaMDS’ function in the R package ‘vegan’^75^, using the Bray-Curtis dissimilarity index computed from relative biovolume data. We then used GAMs to relate these two NMDS axes to glyphosate exposure in Phase I (to determine whether glyphosate forces communities towards a homogeneous composition) and to chlorophyll *a* at the end of Phase II (to determine whether composition predicts rescue). Finally, to further quantify which community variable best predicted rescue in Phase II, we used univariate regression tree analysis and AIC-based model comparison of univariate GAMs. Both analyses used log-transformed chlorophyll *a* at the end of Phase II (‘rescue’) as the response and a number of (scaled) predictor variables hypothesized to influence community response to the lethal dose of glyphosate, namely glyphosate concentrations at the end of Phase I and Phase II (log-transformed), the two NMDS axes, zooplankton density at the end of Phase II, and chlorophyll *a* (log-transformed), genus number, alpha diversity, and the biovolume (log-transformed) of four taxa at the end of Phase I. These taxa were *Selenastrum*, *Ankistrodesmus*, *Desmodesmus*, and *Chlorella*, which collectively accounted for 96.5 % of total biovolume at the end of Phase II (and thus constitute the only taxa that could influence rescue). A conditional inference regression tree with these predictors was fitted with the ‘ctree’ function in the R package ‘party’^76^, using Monte Carlo permutation tests to assess the significance of correlations between each predictor and the response. A separate univariate GAM was also fitted for each of the 13 predictor variables, and model fit (the extent to which each predictor is linked to rescue) was compared with AIC. These two analyses focused on the 16 ponds for which all data requirements were met.

## Data availability

All data presented therein and all computer code used for analyses will be archived on an online repository upon manuscript acceptance.

## Acknowledgements

The Canadian Foundation for Innovation and the Liber Ero Chair in Biodiversity Conservation provided funding to A.G. to construct the LEAP mesocosm facility. The authors also acknowledge support and operating funds from the Natural Sciences and Engineering Research Council of Canada (NSERC), the Fonds de Recherche du Québec – Nature et Technologies (FRQNT), the Canada Research Chair Program (R.D.H.B., A.G., B.J.S.), the Quebec Centre for Biodiversity Science (QCBS), and the Groupe de Recherche Interuniversitaire en Limnologie et environnements aquatiques (GRIL). We also thank David Maneli, Charles Normandin, Alex Arkilanian, and Tara Jagadeesh for assistance in the field, Katherine Velghe for nutrient analyses, Pierre Carrier-Corbeil and Milla Rautio for phytoplankton identification, and Marco Aurelio Piñeda Castro for developing the LC-MS method for glyphosate measurements and for conducting chemical analyses.

## Author contributions

V.F., M.P.H., R.D.H.B., B.E.B, G.B., G.F.F., B.J.S. and A.G. designed the study. V.F., M.P.H., and N.B.C. collected all data. C.C.Y.X. and V.Y. contributed to the development of laboratory methods. B.E.B. provided a trait database. V.F. analyzed data, made the figures, and drafted the manuscript. All authors contributed significantly to data interpretation and commented on manuscript drafts.

